# Chromosome structure due to phospho-mimetic H2A modulates DDR through increased chromatin mobility

**DOI:** 10.1101/2020.11.25.397356

**Authors:** Fabiola Garcia Fernandez, Brenda Lemos, Yasmine Khalil, Renaud Batrin, James E. Haber, Emmanuelle Fabre

## Abstract

In budding yeast and mammals, double strand breaks trigger global chromatin mobility together with the rapid phosphorylation of the histone H2A over an extensive region of the chromatin. To assess the role of H2A phosphorylation in this response to DNA damage, we have constructed strains where H2A has been mutated to the phospho-mimetic H2A-S129E. We show that H2A-S129E mutant increases global motion of chromosomes even in the absence of DNA damage. The intrinsic chromatin mobility of H2A-S129E is not due to checkpoint activation, histone degradation or kinetochore anchoring. Rather, the increased intra-chromosomal distances observed in H2A-S129E mutant are consistent with chromatin structural changes. In this context, the Rad9^53BP1^-dependent-checkpoint becomes dispensable. The increase in chromatin dynamics is favorable to NHEJ of a single double-strand break but is accompanied by a sharp decrease in inter-chromosomal translocation rates. We propose that changes in chromosomal conformation due to H2A phosphorylation are sufficient to modulate the DDR and maintain genome integrity.

## Introduction

Eukaryotic cells have developed sophisticated machineries to respond to the multiple stresses they are constantly confronted with. In the presence of DNA Double-Strand Breaks (DSBs), the DNA damage response (DDR) protects the genome by detecting and repairing the potentially lethal DSBs that could lead to genome instability or tumorigenesis. Inherited or acquired defects in DDR can result in various diseases, such as immune deficiency, neurological degeneration, premature aging, and severe cancer susceptibility (reviewed in (Jackson and Bartek, 2009; Goldstein and Kastan, 2015).

The DDR starts by the recruitment of surveillance proteins that activate cell cycle checkpoint, promote chromatin remodeling allowing DNA accessibility to the repair machinery and trigger DNA repair pathway choice (for reviews see (Mehta and Haber, 2014; Symington and Gautier, 2011; Ceccaldi et al., 2016; Aylon and Kupiec, 2004)). Two of the most well-known DSB repair mechanisms are homologous recombination (HR) and non-homologous end joining (NHEJ). While HR repairs DNA breaks by copying the missing information across the lesion from an undamaged template, as from the replicated sister chromatid, NHEJ does it by ligation of the broken ends after their juxtaposition (reviewed in (Mehta and Haber, 2014; Symington and Gautier, 2011; Ceccaldi et al., 2016; Aylon and Kupiec, 2004)). Long considered as error-prone, the classical form of NHEJ is now regarded as a versatile, adaptable and essential pathway for the maintenance of genomic stability, because joining of the juxtaposed ends of the breaks does not necessarily involve nucleotide deletions. However alternative forms of end-joining exist that can induce DNA aberrations, including chromosomal translocations (reviewed in (Emerson and Bertuch, 2016; Bétermier et al., 2014; McVey and Lee, 2008)).

To repair the damage, the DNA damage checkpoint delays cell cycle progression. This can trigger arrests at the G1/S transition, during the S-phase, or at the G2/M transition, depending on the nature of the damage and the phase of the cell cycle in which the lesion happens (for reviews see (Finn et al., 2011; Shaltiel et al., 2015; Waterman et al., 2020)). The highly conserved MRX^MRN^ complex (Mre11-Rad50-Xrs2 in yeast; Mre11-Rad50-Nbs1 in mammals) is among the earliest sensors of DSB by binding directly to DNA broken ends. Sensing includes activation of the phosphatidylinositol3-kinase-related kinase (PIKK) family such as mammalian ATM (Ataxia-Telangiectasia-Mutated) and ATR (ATM- and Rad3-related), called Tel1 and Mec1, respectively, in budding yeast. These sensing kinases are required independently after a DSB, at different points of the DDR. Tel1^ATM^ is rapidly recruited and activated after the recognition of the DSB by the MRX^MRN^ complex, whereas Mec1^ATR^ is recruited by the ATR-interacting protein Ddc2^ATRIP^ after 5’ to 3’ resection of the DSB ends yields single-stranded DNA and binding of RPA (Kondo et al., 2001; Tibbetts et al., 1999; Nakada et al., 2003; Shroff et al., 2004; Li et al., 2020; Zhou and Elledge, 2000). Both kinases phosphorylate several other proteins involved in cell cycle checkpoint control and DNA repair. An important landmark is the phosphorylation of histone H2A in yeast, or histone variant H2AX in mammals. Phosphorylation of histone H2A occurs near its C terminus (S129 in yeast H2A and S139 in mammalian H2AX, (Rossetto et al., 2014; Redon et al., 2003; Burma et al., 2001; Celeste et al., 2003; Shroff et al., 2004). Phosphorylation of H2A(X) in yeast and mammals, also known as γ-H2A(X), rapidly accumulates at the DSB, spreads over long distances and contributes to further DNA signaling and repair (Downs et al., 2000; Shroff et al., 2004; Burma et al., 2001; Lee et al., 2014; Renkawitz et al., 2013).

In particular, phosphorylation of H2A(X) at the site of DSB allows recruitment of chromatin remodeling complexes (INO80, SWR1) and other proteins, including 53BP1 (the ortholog of yeast Rad9) (Hammet et al., 2007; van Attikum et al., 2004; Morrison et al., 2004; Tsukuda et al., 2005). Rad9^53BP1^ is a critical checkpoint adaptor protein that transmits the signal from Mec1^ATM^ and Tel1^ATR^ to the downstream effector Rad53^CHK2^ and Chk1^CHK1^ (Blankley and Lydall, 2004; Harrison and Haber, 2006; Sweeney et al., 2005). Rad9^53BP1^ was originally identified in a pioneering study in budding yeast because it controls cell cycle progression by arresting cells in the G2/M phase in case of unrepaired damage (Weinert and Hartwell, 1988). Rad9 and 53BP1 both contain BRCT and Tudor domains that recognize histone H3 methylation as well as γ-H2A(X) phosphorylation (Hammet et al., 2007; Lancelot et al., 2007). Deletion of Rad9 or 53BP1 results in a more rapid degradation of DSB ends (Lazzaro et al., 2008; Ferrari et al., 2015) Rad9^53BP1^’s protective role in genome integrity is evidenced by Δ*rad9*’s poor survival upon genotoxic treatments including phleomycin, γ-rays and UV (Menin et al., 2019; Mirman and de Lange, 2020)

Neither the functional relevance of chromatin modifications nor the mechanism by which the DDR is activated by chromatin itself is fully understood. It has been shown that robust targeting of repair factors or kinase sensors, in the absence of damage, can elicit the DDR both in yeast or in mammals, indicating that local concentration of sensor protein and/or the higher order of chromatin structure are key in the DDR cascade (Soutoglou and Misteli, 2008; Bonilla et al., 2008). In fact, in mammals, chromatin at the site of damage experiences two successive waves of changes. A first local expansion dependent on γ-H2A(X) and MRN(X) is followed by a dynamic compaction, the latter being enough to activate the DDR (Ziv et al., 2006; Kruhlak et al., 2006; Khurana et al., 2014; Burgess et al., 2014). Chromatin compaction, due to the tethering of methyltransferase SUV3-9 or heterochromatin protein HP1, can trigger ATM signaling and activate upstream (γ-H2A(X)) but not downstream (53BP1) components of the DDR cascade, showing that chromatin compaction is an integral step of the DDR (Burgess et al., 2014). Strikingly, a global alteration of chromatin and chromosome structure, induced by hypotonic condition or mechanical stress, is sufficient to activate ATM or ATR in the absence of any DNA damage, respectively, (Bakkenist and Kastan, 2003; Kumar et al., 2014). Yeast lacks homologs of SUV3-9 or HP1, but interestingly, Mec1^ATR^ might respond to chromatin dynamics during S phase, when replication stress occurs and causes mechanical stress on the nuclear membrane (Forey et al., 2020; Bermejo et al., 2011).

Global modification of damaged genomes is also evidenced through an increase of chromosome mobility, a process that may favor repair in yeast (Miné-Hattab and Rothstein, 2012; Seeber et al., 2013; Strecker et al., 2016; Hauer et al., 2017; Herbert et al., 2017; Lawrimore et al., 2017; Smith et al., 2018). The molecular mechanisms at play in global mobility are not fully deciphered, but regulatory networks are becoming clearer (reviewed in (Zimmer and Fabre, 2018; Smith and Rothstein, 2017; Seeber et al., 2018; Haber, 2018)). The DDR activation by Mec1^ATM^ and Tel1^ATR^ is a critical first step in the response (Seeber et al., 2013; Smith et al., 2018; Dion et al., 2012; Miné-Hattab and Rothstein, 2012). Mec1^ATM^ and Tel1^ATR^ activation have diverse consequences, such as centromeric relaxation (Strecker:2016jj; Lawrimore et al., 2017) and chromatin modifications (Herbert et al., 2017; Hauer et al., 2017; Miné-Hattab et al., 2017). It has been proposed that the repulsive forces of the negative charges due to H2A phosphorylation could render the chromosome stiffer (Herbert et al., 2017), or that histone depletion could be the cause of a more expanded chromatin (Hauer et al., 2017; Cheblal et al., 2020), both chromatin alterations resulting in the observed increase of chromosome mobility. It is remarkable that increased chromosome mobility, as a response to genomic insults, is a conserved feature in metazoan genomes, with regulations involving repair proteins 53BP1 or γ-H2AX, like in yeast (Ryu et al., 2015; Clouaire et al., 2018; Lottersberger et al., 2015; Dimitrova et al., 2008; Schrank et al., 2018).

To better understand how γ-H2A(X) affects different aspects of chromosome structure and movement, we have created yeast strains in which both copies of the H2A encoding genes have been mutated to S129E, which appears to be phosphomimetic (Eapen et al., 2012). We find that H2A-S129E fully recapitulates the increased apparent stiffness of chromatin observed after damage, resulting in increased intra-chromosomal distances and chromosome dynamics. An increased repair by NHEJ and a decreased rate of translocation correlate with this change in chromatin. Surprisingly, we find that H2A-S129E rescues Δ*rad9* survival deficiency after DNA damage, further supporting the notion that changes in chromatin structure are a key contributor to DNA damage response.

## Results

### In the absence of DNA damage, mimicking H2A phosphorylation increases chromosome mobility

To study the consequences of mimicking H2A-S129 phosphorylation on chromosome mobility, we created isogenic yeast strains in which both *HTA1* and *HTA2* were mutated to H2A-S129E by CRISPR/Cas9-mediated gene editing (see Materials and Methods). We used four strains (P1-P4) that carry fluorescently labeled green loci (LacI-GFP bound to an array of LacO sites) at different regions of the right arm of chromosome IV (**Figure 1A**). As control, we replaced S129 by an alanine to impede phosphorylation, using the same strategy (H2A-S129A). We first compared the growth of the H2A-S129 mutants to their wild-type counterpart, in the presence or in the absence of DNA damage generated by the radio-mimetic drug Zeocin (Dion et al., 2012; Seeber et al., 2013; Herbert et al., 2017). In the absence of Zeocin, growth rates of P1-P4 WT, H2A-S129E and H2A-S129A strains were indistinguishable, suggesting no particular endogenous damage caused by these mutations. WT strains exhibited a sensitivity to Zeocin (**Figure 1A, Supplementary Figure 1A**), as previously shown (Seeber et al., 2013; Strecker et al., 2016; Krol et al., 2015). In the presence of Zeocin, growth of H2A-S129E was similar to that of WT, whereas H2A-S129A mutants showed increased sensitivity (**Figure 1A, Supplementary Figure 1A**). To precisely measure the sensitivity to Zeocin of the mutant strains, we performed colony forming unit (CFU) assays by calculating the ratio between colonies grown on media with and without Zeocin. CFU ratios were close to 70% in P1-P4 WT cells (**Figure 1B**). Likewise, CFU ratios in H2A-S129E mutant strains ranged between 81% ± 3.1 and 96 ± 3.2%, with no significant difference from WT, except for the P4 strain, which showed a slightly higher survival (Downs et al., 2000; Moore et al., 2007; Redon et al., 2003). In H2A-S129A strains, CFU ranged from 47.5% ± 4.0 to 64.5% ± 7.2 confirming the enhanced sensitivity of these mutants to Zeocin compared to WT, as previously shown (Downs et al., 2000; Moore et al., 2007; Redon et al., 2003). The simplest interpretation for a similar survival to DNA damage between H2A-S129E mutant and WT is that glutamic acid can replace the function normally provided by S129 phosphorylation, a conclusion further supported by the analysis of the H2A-S129A mutants.

**Figure 1.**
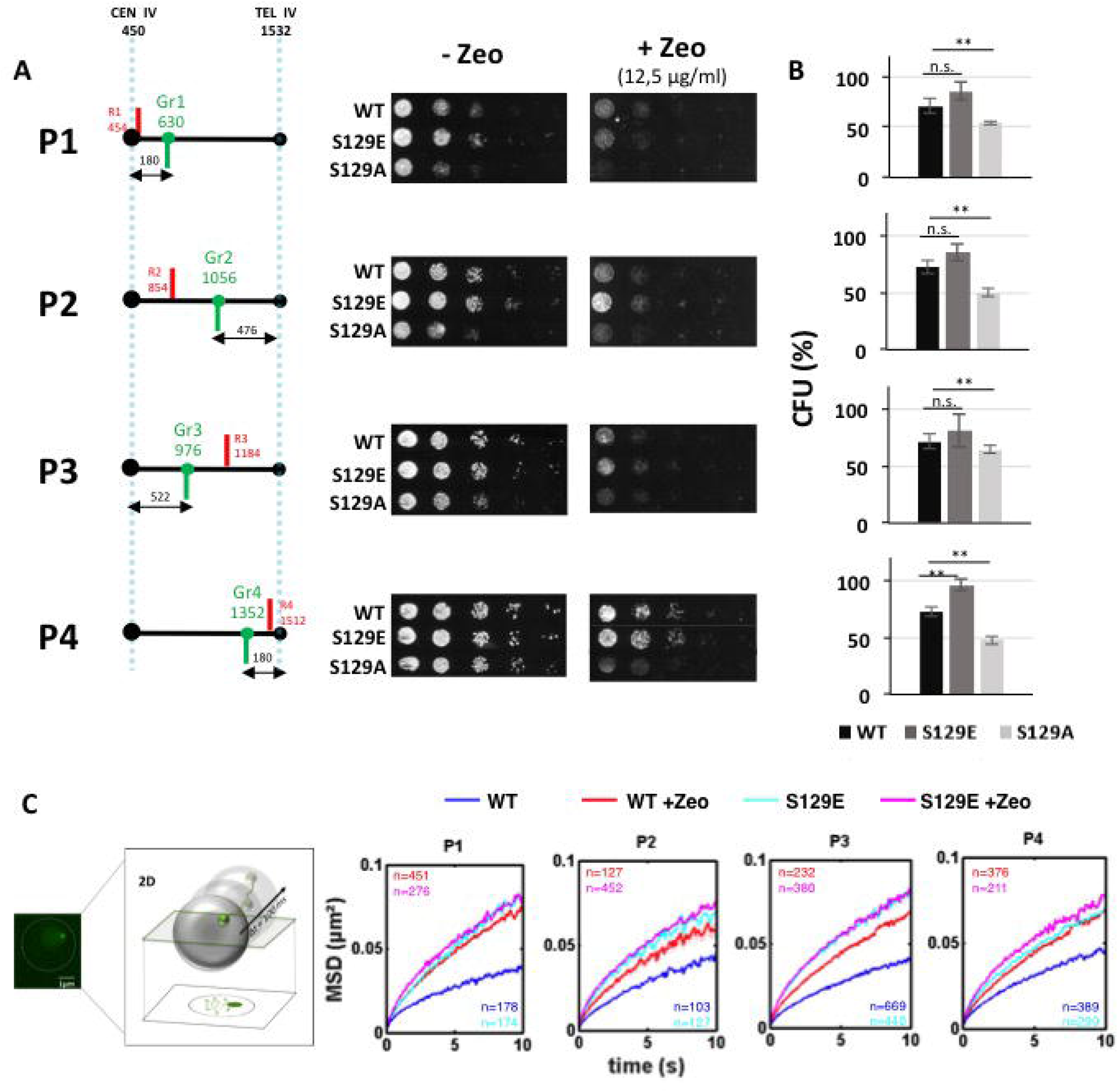
H2A-S129E increases global dynamics in absence of DNA damage. **A.** Left, schematics of chromosome IV, indicating the genomic positions of fluorescently labeled loci investigated in this study. Numbers above the black line indicate genomic distance from the centromere in kb. Blue dashed bars indicate the centromere (CEN IV) and the right telomere (TEL IVR). Red and green bars indicate the four loci tagged in red (R1– R4) or in green (Gr1-Gr4). Strains P1 to P4 correspond to individual red-green pairs, separated by ~200kb (R1-Gr1, R2-Gr2, Gr3-R3, Gr4-R4, respectively). Right, drop assays (tenfold dilutions) showing comparable growth of wild-type (WT) and H2A-S129E mutants in the absence of damage and comparable sensitivity to long exposition to Zeocin (12, 5μg/ml). As a control, effect of the Zeocin on cell survival is shown for H2A-S129A mutants. **B**. Colony forming units (CFU) calculated as the ratio of cells growing in the presence Zeocin over cells grown in the absence of Zeocin after spreading of ~ 200 colonies on each medium, for the four strains tested (P1-P4) in wild-type (WT, black bars), H2A-S129E mutants (grey bars) and H2A-S129A mutants (light grey bars). Error bars represent the standard deviation of the mean CFU counted of at least three independent experiments; P values are calculated after a non-parametric t test (n.s, not significant P>0.05, * P≤0.05, ** P≤0.01, *** P≤0.0001). **C**. Mean square displacements (MSDs) of Wild-Type cells and H2A-S129E mutant as function of time interval of the four Gr1 to Gr4 green loci in P1 to P4 strains, as computed from 2D time-lapse microscopy data (cell population average). Blue and red curves are for untreated and Zeocin-treated wild-type cells (WT) respectively; cyan and magenta curves are for untreated and treated H2A-S129E mutant, respectively. The numbers of cells used to compute each curve (n) are indicated.

We then explored chromosome mobility in H2A-S129E mutant cells by tracking the fluorescently labeled green loci (Herbert et al., 2017). Green labels are located at 180 kb and 522 kb from the centromere IV or at 476kb and 180kb from the right telomere IV (Gr1, Gr3, Gr2 and Gr4 respectively, **Figure 1A**). We analyzed Mean Square Displacements (MSDs) of the four Gr1-Gr4 loci by tracking hundreds of cells in each P1-P4 strain, by high-speed 100ms time-lapse microscopy, for a time period of 5 minutes (Spichal et al., 2016; Herbert et al., 2017). To induce DNA damage, we used a final concentration of Zeocin of 250 μg/ml for 6h. Under these conditions, in which ~ 80% of the cell population showed damage, as seen by Rad52-GFP foci formation (**Supplementary Figure 1**) (Herbert et al., 2017), WT cells exhibited a global increase in chromosome mobility (**Figure 1C**). Strikingly, in the absence of Zeocin treatment, the MSDs in the H2A-S129E mutant strains, were as high as those observed in wild type strains after induction of damage (**Figure 1C** and **Supplementary Figure 2A**). In these mutant cells, addition of Zeocin for 6h did not significantly increase MSDs, suggesting that H2A-S129E mutation maximizes mobility that Zeocin cannot increase any further (**Figure 1C and Supplementary Figure 2A**). After Zeocin exposure, the lower increase in mobility of the four tagged Gr1-Gr4 loci in H2A-S129A mutated cells (**Supplementary Figure 2B** and **2C**), confirmed that γ-H2A(X) is required for full global mobility upon DNA damage (Herbert et al., 2017). Together, these results suggest that the function normally performed by H2A phosphorylation is mirrored by H2A-S129E and that this mutation is sufficient to induce an increase in chromosomal dynamics.

### H2A-S129E mutant does not trigger cell cycle arrest, histone loss or centromere detachment

To understand the mechanism underlying the enhanced dynamics observed in the H2A-S129E mutant, we explored different hypotheses. First, it is well documented that checkpoint activation induces global mobility upon DNA damage (Seeber et al., 2013; Hauer et al., 2017; Smith et al., 2018). To test whether H2A-S129E may induce mobility due to the physiological DDR, we explored the effect of this mutant on cell cycle progression. FACS results on asynchronous populations indicated an equally low number (19-23%) of cells in the G2/M cell cycle phase in the WT and H2A-S129E mutant strains, indicating that the DDR checkpoint leading to cell cycle arrest is not activated in H2A-S129E (**Figure 2A**). In addition, phosphorylation of Rad53, the signal for effective checkpoint (Pellicioli et al., 2001), was not observed in both strains in the absence of damage (**Figure 2A**). However, both WT and H2A-S129E mutant strains exhibited a similar DNA damage response after 6h Zeocin treatment, since the percentage of G2/M arrested cells was enriched (71-77%) in both strains, with an accompanying hyper-phosphorylation of Rad53 (**Figure 2A, Supplementary Figure 3**). These results reject the hypothesis that enhanced chromatin dynamics observed in H2A-S129E mutants in the absence of damage is triggered by constitutive activation of the cell cycle checkpoint. Second, histone loss was shown to elicit enhanced chromatin mobility (Hauer et al., 2017). We therefore examined the global stability of histones. By immuno-blotting, we found comparable levels of total H4 in WT and H2A-S129E mutant cells (**Figure 2B**). Total H4 abundance strongly decreased upon 6h of Zeocin treatment, as previously reported (**Figure 2B**, (Hauer et al., 2017)). Finally a previous study had shown that induction of a single DSB in the genome led to kinetochore protein Cep3 phosphorylation by Mec1 and also correlated with an increase in global mobility associated with an increase in Spindle Pole Body (SPB) - Centromere (CEN) 2D distances (Strecker et al., 2016). We therefore measured the distances between SPC42, a protein of the SPB fused to mCherry and CenIV labeled by an array of TetO repeats bound by tetR-GFP (He et al., 2000). In the absence of Zeocin, the SPB-CenIV distances between WT or H2A-S129E strains were identical and in both cases the distances increased after cells were treated by Zeocin (**Figure 2C**), indicating that a general loosening of the anchoring of the centromeres to the SPB is probably not the cause of the increase in chromosomal mobility observed in H2A-S129E mutants. Thus, the high intrinsic mobility of H2A-S129E mutated chromatin must be due to factors other than checkpoint activation, histone degradation or loosening of centromere tethering to the spindle pole body.

**Figure 2.**
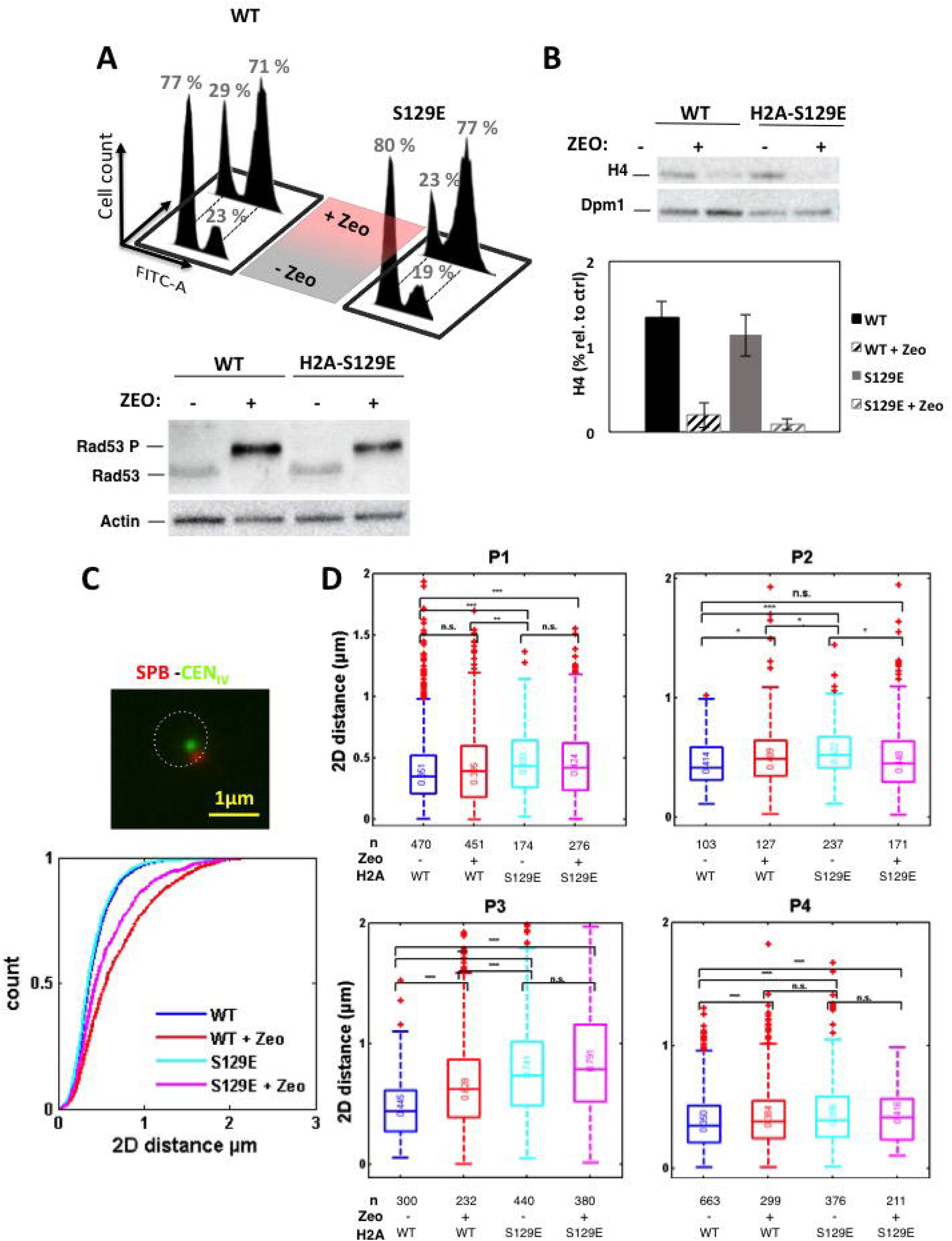
Increase in global dynamics in H2A-S129 mutated cells is not linked to checkpoint activation, histone loss or defective centromeric tethering but to increased intra-chromosomal distances. **A.** Flow cytometry analyses of asynchronous WT and H2A-S129E mutant cell populations treated or not by Zeocin. H2A-S129E mutants show normal cell cycle in the absence of damage and a G2/M arrest in the presence of damage, similarly to the WT. Statistical analyses between WT and H2A-S129E mutated cells with a chi square-test shows not significative values of 0.32 and 0.15 for untreated and treated conditions, respectively. Representative results from immunoblotting showing the phosphorylation status of Rad53–HA in WT and H2A-S129E cells in response or not to Zeocin treatment. Actin was used as a loading control. **B.** Immunoblot analyses and quantification with H4 specific antibodies on whole cell extracts in WT and H2A-S129E mutant before and after Zeocin treatment. Dpm1 was used as a loading control. Immunoblot quantification with ImageJ (mean and s.d.). **C.** Cumulative distributive function (CDF) from SPB-CenIV 2D distances of WT and H2A-S129E mutant cell populations treated or not with Zeocin. **D.** Bar plots show 2D distances of untreated WT (blue) and H2A-S129E (cyan) and Zeocin-treated (red and magenta, respectively), for the four pairs of green-red labeled loci in P1-P4 strains. The horizontal line at the center of each box indicates the median value; the bottom and top limits indicate the lower and upper quartiles, respectively. The whiskers indicate the full range of measured values, except for outliers, which are shown as small red dots. Brackets indicate the result of a Wilcoxon rank-sum test between distributions with “n.s.” for “not significant” for P > 0.05, * for P ≤ 0.05, ** for P ≤ 10^−2^ and *** for P ≤ 10^−3^. Number of analyzed (n) cells is indicated.

### Mimicking H2A-S129 phosphorylation increases intra-chromosomal distances without DNA damage

It has been established that an increase in genomic mobility could be explained by a change in chromatin structure (Hauer et al., 2017; Herbert et al., 2017; Miné-Hattab et al., 2017). Based on polymer modeling and multi-scale tracking of chromatin after damage, a global stiffening of the chromatin fiber is consistent with a simultaneous increase in chromosomal mobility and spatial distances between loci on the same chromosome (Herbert et al., 2017; Miné-Hattab et al., 2017). H2A phosphorylation was proposed as a potential molecular mechanism, since the negative repulsive charges due to H2A phosphorylation could increase the stiffness of the chromatin fiber, as seen *in vitro* and by modeling (Cui and Bustamante, 2000; Qian et al., 2013; Herbert et al., 2017). In addition, based on similar experiments, but using different modeling assumptions, chromatin decompaction has also been proposed to play a role (Hauer et al., 2017; Amitai et al., 2017). We therefore hypothesized that the increase in H2A-S129E chromosome motion could be linked to a change in chromosomal structure. To address this question, we measured red-green pairwise distances in P1-P4 WT and H2A-S129E mutated strains (**Figure 2D**). In these strains, TetO arrays bound by tetR-mRFP are inserted at about 200 kb from the green labels (see **Figure 1A**). As expected, intra-chromosomal distances increased in WT strains after prolonged treatment to Zeocin (3h), ranging from (~ 350 - 414nm to ~ 384 - 628nm in the absence and presence of Zeocin respectively, (**Figure 2D**, (Herbert et al., 2017)). In the absence of Zeocin treatment, large increases of intra-chromosomal distances (from ~ 396 - 496 nm) were observed all along the chromosome arm in H2A-S129E mutants, similar to those observed in WT upon Zeocin treatment (**Figure 2D**). In contrast, intra-chromosomal distances do not increase in H2A-S129A mutants (Herbert et al., 2017), showing that the effect on distances was specific to H2A-S129E mutation. A structural change in chromatin is accordingly the most probable cause for the increased mobility observed in the H2A-S129E mutant, in the absence of any DNA damage.

### Rad9 checkpoint control on cell survival and global mobility upon Zeocin treatment is suppressed by mimicking H2A phosphorylation

The observation that H2A-S129E causes an increase in chromosome dynamics independent of cell cycle arrest suggests that structural changes in the chromatin of this mutant may make checkpoint factors dispensable for damaged-induced chromatin mobility (Seeber et al., 2013; Smith et al., 2018; Dion et al., 2012; Miné-Hattab and Rothstein, 2012; Bonilla et al., 2008). To test this hypothesis, we mutated the key checkpoint adaptor Rad9^53BP1^ protein in the DDR that mediates the interaction between modified histones such as γ-H2A(X) and several effector proteins in the DDR (Finn et al., 2011). We deleted *RAD9* in the WT and H2A-S129E P2 strains and analyzed single (Δ*rad9*) and double (Δ*rad9* H2A-S129E) mutant phenotypes on cell survival, cell cycle profile and chromosome mobility. In the absence of Zeocin treatment, both mutants grew similarly to their wild type *RAD9* counterpart in spot assays (**Figure 3A**). As expected, the growth of the Δ*rad9* mutant was affected in the presence of Zeocin (Seeber et al., 2013). Interestingly, spot assay and CFU ratio quantification of treated versus non-treated cells, indicated that the Δ*rad9* H2A-S129E double mutant was indistinguishable from WT or H2A-S129E single mutant strains (**Figure 3A, 3B**). Moreover, while cell cycle arrest due to Zeocin was deficient in Δ*rad9* mutant, as predicted, it was restored in Δ*rad9* H2A-S129E double mutant (after Zeocin treatment, p=0.49 between WT and Δrad9 H2A-S129E double mutant and p=4.4 10^−14^ between WT and Δrad9, as determined by a contingency chi-square test (**Figures 2A, 3C**). Surprisingly, Rad53 phosphorylation was not restored in the Δ*rad9*, H2A-S129E double mutant (**Figures 3C**). Thus, the failure of the Δ*rad9* mutant to survive and activate cell cycle arrest in response to Zeocin, is rescued by H2A-S129E mutation independently of Rad53 phosphorylation. These results suggest that an early DDR response – the phosphorylation of histone H2A – enables cells to delay cell division in the absence of Rad9 and thus provide more time for DNA repair (see Discussion).

**Figure 3.**
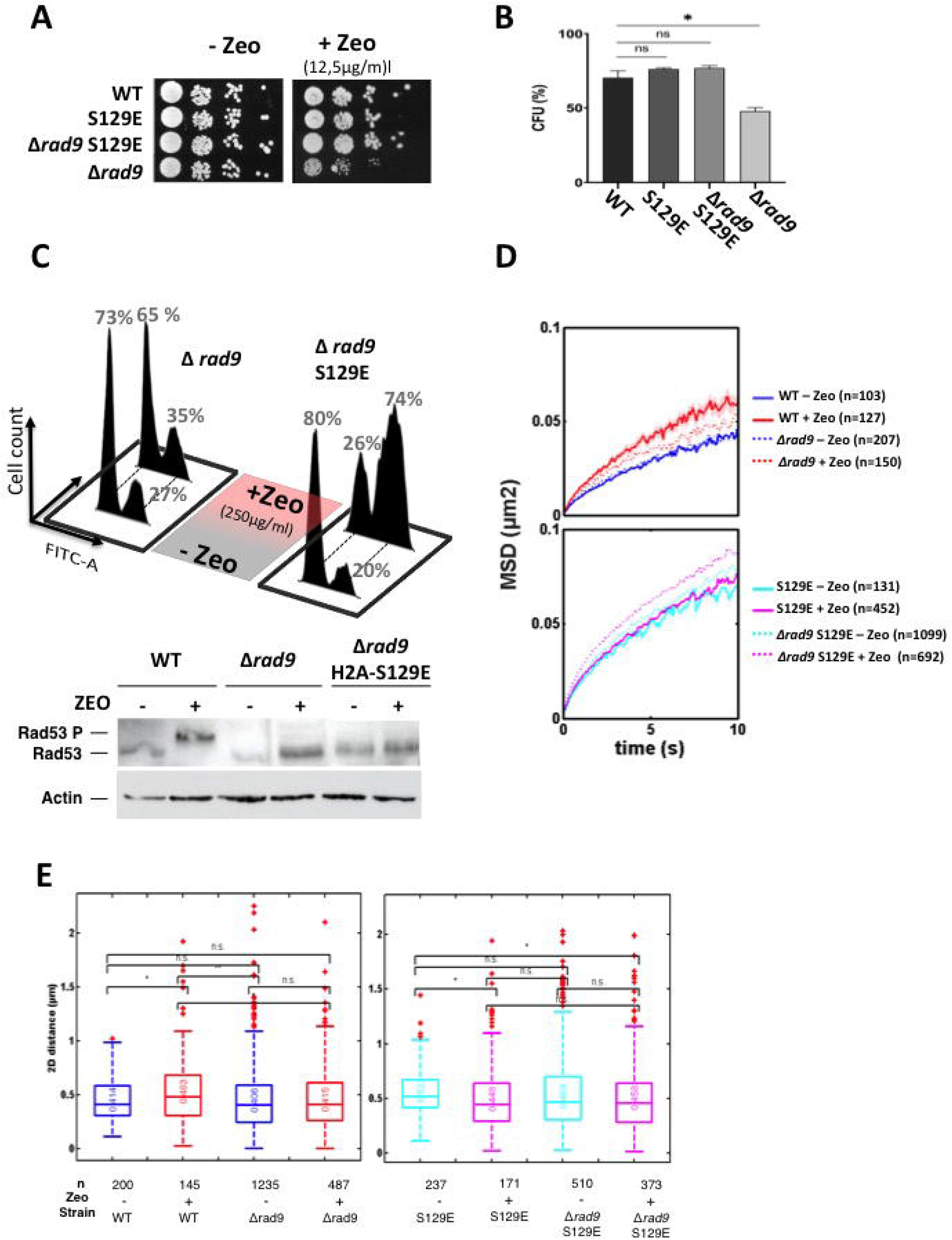
H2A-S129E mutant suppresses Rad9 checkpoint control on cell survival, global dynamics and intra-chromosomal distances upon Zeocin treatment. **A.** Drop assay (tenfold dilutions) of Wild-Type (WT), Δ*rad9* and Δ*rad9* H2A-S129E double mutant in the absence and the presence of 12,5μg/ml of Zeocin for the P3 strain. The single H2A-S129E mutant is also shown. **B**. Colony forming units (CFU) calculated as the ratio of cells growing in the presence Zeocin over cells grown in the absence of Zeocin after spreading of ~ 200 colonies on each medium in wild-type (WT, black bar), H2A-S129E mutant (dark grey bar), Δ*rad9* (light grey bar) and Δ*rad9* H2A-S129E double mutant (grey bar). Error bars represent the standard deviation of the mean CFU counted of at least three independent experiments; P values are calculated after a not-parametric t test (n.s, not-significant P>0.05, * P≤0.05, ** P≤0.01, *** P≤0.0001). **C.** FACS analyses in the same strains in the absence or the presence of 250μg/ml of Zeocin. Representative results from immunoblotting showing the phosphorylation status of Rad53–HA in WT, Δ*rad9* and Δ*rad9*, H2A-S129E cells in response or not to Zeocin treatment. Actin was used as a loading control. **D.** Mean square displacements (MSDs) calculated as in figure 1, WT compared to Δrad9 on the top; H2A-S129E compared to Δ*rad9* H2A-S129E double mutant on the bottom. Non-treated cells are blue, dotted blue, cyan and dotted cyan and treated cells are red, dotted red, magenta and dotted magenta, respectively. **E** Box plots show 2D distances of untreated and Zeocin-treated cells (same color code as in **D**) of WT compared to Δ*rad9* on the left and H2A-S129E compared to the double mutant Δ*rad9* H2A-S129E on the right. Analyzed cells range from (n) ~150 - ~1200. P values are calculated after a non-parametric t test (n.s, not significant P>0.05, * P≤0.05, ** P≤0.01, *** P≤0.0001).

We then measured global chromatin mobility in these mutants in the absence and the presence of Zeocin (**Figure 3D**). Whereas the Δ*rad9* mutant behaved similarly to WT in undamaged conditions, the absence of *RAD9* partially impeded the enhancement of global chromatin mobility observed in WT strains upon DNA damage, in agreement with (Seeber et al., 2013). Interestingly, in both undamaged and damaged conditions, the Δ*rad9* H2A-S129E double mutant induced a massive increase of global dynamics, comparable to the single H2A-S129E mutant (**Figure 3D, Supplementary Figure 4**). To check whether changes in chromosome dynamics also translate into chromatin structural changes in the Δ*rad9* H2A-S129E double mutant, we also measured intra-chromosomal 2D distances (**Figure 3E**). We confirmed that in the absence of Zeocin treatment, the distances in Δ*rad9* mutant were not significantly different to those of the wild-type strain and that the distances in the Δ*rad9* H2A-S129E double mutant were comparable to the single H2A-S129E mutant. As expected from the dynamic behavior, in the presence of Zeocin, Δ*rad9* showed a modest increase in intra-chromosomal distances as compared to the WT but increase as much as H2A-S129E in the double Δ*rad9* H2A-S129E mutant (**Figure 3E**). Thus, the structural changes induced by the H2A-S129E mutant in the Δ*rad9* background corroborate the observations on the chromosomal dynamics. Our results highlight the capacity of H2A-S129E mutant to suppress the specific contribution of Rad9 to cell survival, DNA damage checkpoint and global chromosome dynamics when genome is damaged, possibly through chromatin structure modification.

### Mimicking H2A-S129 phosphorylation increases NHEJ and reduces translocation rates of Δrad9

Rad9 and its mammalian ortholog 53BP1, are known to facilitate NHEJ (Ferrari et al., 2015; Zimmermann et al., 2013). Since the Zeocin sensitivity of Δ*rad9* is suppressed by the H2A-S19E mutant that increases chromosome mobility, and since chromosome mobility could be a means for promoting the joining of double-stranded extremities by NHEJ (Lottersberger et al., 2015; Dimitrova et al., 2008), we investigated the effect of both Δ*rad9* and H2AS129E mutants on the NHEJ repair pathway. We used a plasmid repair assay, in which repair of a linear plasmid can only be mediated by NHEJ. We generated linear plasmids with cohesive ends, by cutting a *HIS3*-containing centromeric plasmid by *EcoRI* enzyme. In this assay, only circularized plasmids confer histidine prototrophy to the transformed cells. A Δ*yku70* mutant, which is impaired in NHEJ, was used as control for plasmid linearization. After transformation by the digested plasmids, His^+^ colonies were obtained in the Δ*yku70* mutant 10-fold less than in WT cells. These data indicate that most transforming molecules were efficiently re-circularized in WT (**Figure 4A**). The number of His^+^ colonies recovered in the H2A-S129A mutant was almost as low as in Δ*yku70* mutant, revealing that plasmid circularization by NHEJ is compromised in the absence of γ-H2A(X). This result is consistent with previous studies showing that truncated forms of H2A lacking S129 are deficient for NHEJ and is consistent with an increased resection rate observed in H2A-S129A mutant (Downs et al., 2000; Moore et al., 2007; Eapen et al., 2012). In contrast, the number of His^+^ colonies was ~ 1.3 times higher in the H2A-S129E mutant than in WT, indicating that mimicking H2A phosphorylation allowed more effective repair by NHEJ than in WT (**Figure 4B**). In the Δ*rad9* mutant, the number of His^+^ cells was 4 to 8 times lower than in the WT (**Figure 4B**), an expected result given the role of Rad9 in limiting 5’ DNA end resection (Ferrari et al., 2015). Remarkably, the number of His^+^ colonies was similar to WT in the Δ*rad9*, H2A-S129E double mutant. Thus the restoration of cell survival to DNA damage observed in Δ*rad9* H2A-S129E double mutant could be linked to an increase in NHEJ efficiency. In turn, increased chromosome motion observed in H2A-S129E could promote NHEJ, as it was observed for dysfunctional telomeres in mammalian cells (Lottersberger et al., 2015; Dimitrova et al., 2008).

**Figure 4.**
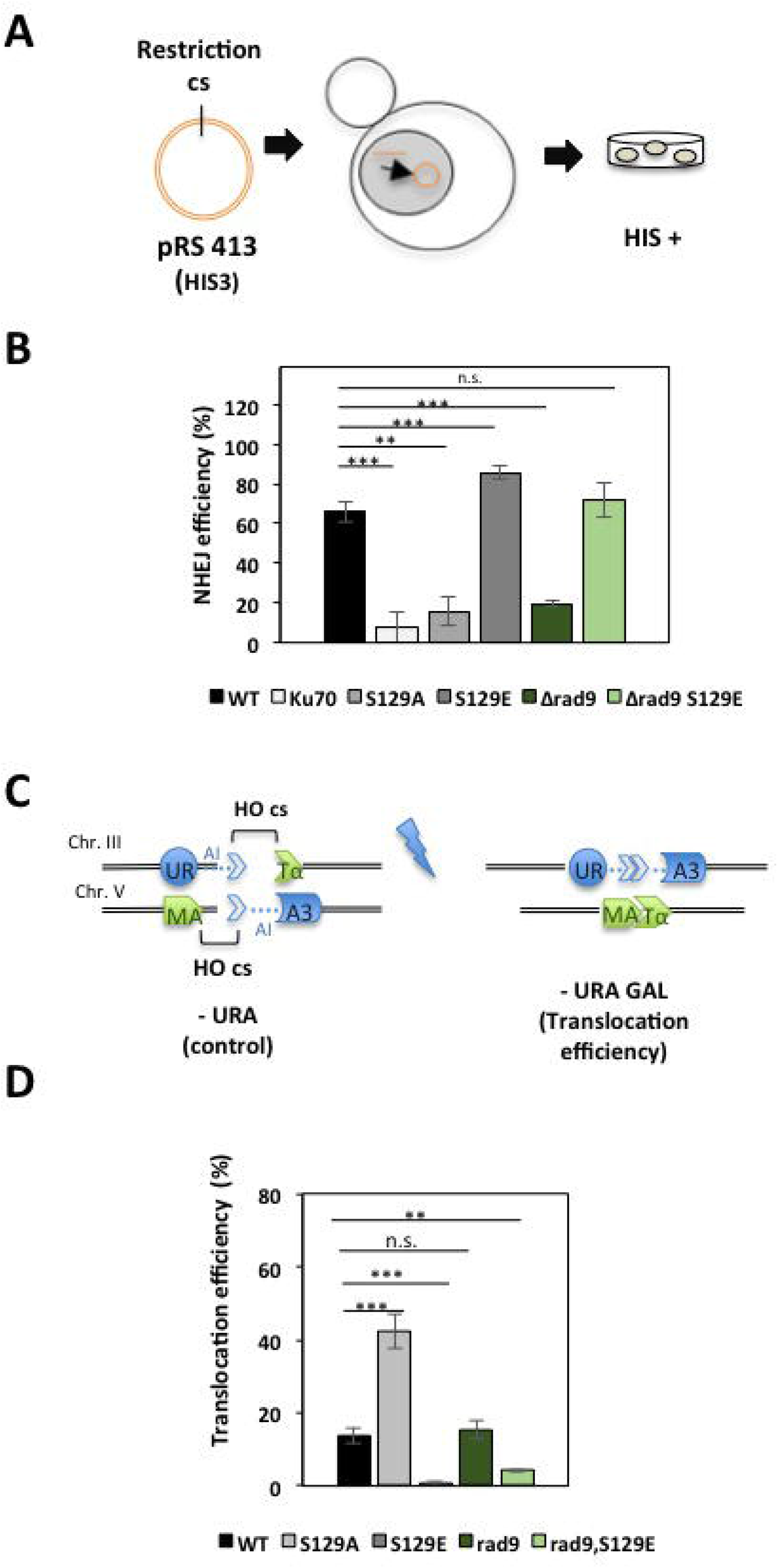
Mimicking H2A S129 phosphorylation increases NHEJ and decreases translocation rates. **A.** Schematic of the NHEJ assay principle. A replicative plasmid carrying HIS3 as an auxotrophic marker (pRS413) is linearized by *EcoR*I enzyme and used for yeast transformation. In strains efficient for NHEJ, plasmid extremities are joined and His+ colonies are recovered on plates lacking histidine (-HIS). **B.** Rejoining efficiency is calculated as the ratio of His^+^ colonies recovered after transformation of a linearized plasmid relative to introduction of a non-linearized plasmid. A Δ*yku70* mutant is impaired in NHEJ and is used as a control for plasmid digestion efficiency. Bar graphs show mean ± s.e.m. P values are calculated after a non-parametric t test (n.s, not significant P>0.05, * P≤0.05, ** P≤0.01, *** P≤0.0001). **C**. Schematic of the strain used to measure translocation. Two DSBs generated by HO endonuclease (HOcs), an intronic sequence and truncated forms of *URA3* are depicted on two distinct chromosomes. Reciprocal end-joining produces Ura^+^ cells. **D.** Translocation efficiency is calculated as the ratio of Ura^+^ colonies recovered after DSB persistent induction (-URA plates containing Galactose) relative to non-translocated conditions (SC plates containing Galactose). P values are calculated after a non-parametric t test (n.s, not significant P>0.05, * P≤0.05, ** P≤0.01, *** P≤0.0001).

We next explored whether large-scale chromatin mobility could also promote the joining of DSBs that are at long range distance, leading to translocation mis-repair, as observed in mammals (Lottersberger et al., 2015). To test for translocation rates, we used a genetic system in which recovery of Ura^+^ *MATα* colonies is mediated by a reciprocal translocation between two DSBs generated in truncated *URA3* on two distinct chromosomes, chromosomes III and V (Lee et al., 2008) (**Figure 4C**). DSBs were induced by expressing the HO endonuclease from the galactose inducible *GAL1-10* promoter. We first checked by qPCR that the cutting efficiency was similar after 1h of galactose induction at the two HO cutting sites in WT and H2A-S129E mutant cells (**Supplementary Figure 5**). Then, translocation rates were measured after permanent induction of HO on galactose-containing plates. The number of Ura^+^ colonies growing on GAL-URA plates was divided by the total number of cells grown in non-selective YPGAL plates for each strain. While phospho-deficient H2A-S129A mutant increased translocation rates ~ 3-fold compared to WT (13.5% ± 2.6 and 42.4% ± 4.8 for WT and H2A-S129A, respectively), as previously documented (**Figure 4D** and (Lee et al., 2008)), H2A-S129E decreased translocation rates by approximately 13-fold (1.1% ± 0.6%) (**Figure 4D**). As expected, the efficiency of translocations in the mutant Δ*rad9* is comparable to that of WT (**Figure 4D** and (Lee et al., 2008)) and significantly reduced in the double mutant Δ*rad9* H2A-S129E (**Figure 4D**).

Altogether, our results indicate that H2A phosphorylation mimicry induces an increase in chromosomal dynamics favorable to a local NHEJ repair concomitant with a sharp decrease in the inter-chromosomal translocation rate.

## Discussion

Here we show that in the absence of DNA damage induction, the H2A-S129E mutation fully recapitulates global chromosome/chromatin dynamics previously observed upon Zeocin treatment (Hauer et al., 2017; Herbert et al., 2017). The specificity of this H2A-S129E mutant is to mimic H2A phosphorylation by the criterion of being recognized by a γ-H2AX-specific antibody (Eapen et al., 2012). One intriguing consequence of H2A-S129E mutation is to allow bypassing an essential function of the DNA damage checkpoint protein, *RAD9*^53BP1^, when Zeocin is present. Locus tracking in living cells reveals that H2A-S129E mutant restores the high chromatin mobility induced by DNA damage, which is defective in Δ*rad9*. Enhanced global dynamics was not related to tethering of the centromeres, histone depletion or checkpoint activation, these mechanisms being proposed to explain the increase in overall chromosome mobility observed when the genome is damaged (Strecker et al., 2016; Smith et al., 2018; Lawrimore et al., 2017; Seeber et al., 2013). Rather, we found that the negative charges of glutamic acid (but not the uncharged alanine) enhanced global dynamics together with increased intra-chromosomal distances. These two characteristics indicate a structural modification of the chromatin compatible with its stiffening (Herbert et al., 2017). If transcription of some key regulators involved in the DDR would be modified in H2A-S129E mutated cells, a similar phenotype could be expected. However, no such changes in transcription could be detected by RNAseq analyses of H2A-S129E mutant compared to WT (S. Bohn, B. Lemos, J.E. Haber and N. Krogan, unpublished result). The fact that Δ*rad9* DNA damage sensitivity and checkpoint defects are rescued by H2A-S129E suggests that H2A-S129E may have a positive effect on repair. We tested this proposal by examining the repair by NHEJ of a linearized plasmid and found that it was favored in H2A-S129E cells. These observations are consistent with the idea that chromatin mobility of DNA extremities may favor their rejoining as proposed in mammalian cells (Lottersberger et al., 2015). However, the direct measurement of DSB translocations revealed that the level of joining in *trans* was severely reduced in H2A-S129E. We propose that changes in chromosomal conformation due to H2A phosphorylation are a means to efficiently modulate the DDR.

### A model for the regulation of DDR by changes in chromatin structure

Chromatin structure is one of the key factors in the DDR since it is the first in being altered both by the damage itself that generates DSBs and by the accumulation of histone modifications and repair proteins at the damaged site. DSB might alter topological constraints on the chromatin fiber, although little is known about these constraints associated with higher order chromosome organization (Canela et al., 2017). Histone post-translational modifications on the other hand, such as phosphorylation of H2A, which can extend from the DSB over areas as large as 50-kb on either side of the DSB (Shroff et al., 2004; Lee et al., 2014), also change the chromatin structure, the repulsive negative charges due to phosphorylation being compatible with a stiffer chromatin (Reina-San-Martin et al., 2003; Celeste et al., 2003; Lee et al., 2008; Miné-Hattab et al., 2017; Herbert et al., 2017). Normally, γ-H2AX modifications occur around the site of a DSB, which exhibits enhanced local motion, while inducing global mobility changes in undamaged regions. Here we see that H2A-S129E-containing chromatin in undamaged regions display enhanced mobility. Whether this represents global motion or is a manifestation of a local motion, we yet cannot determine.

In mammals, regulation of ATM response by chromatin structure was also proposed to contribute to the extremely rapid and sensitive response to damage, across the nucleus, as shown by ATM activation in the absence of detectable damage, during hypotonic shock or during histone deacetylation (Bakkenist and Kastan, 2003; Kumar et al., 2014). The changes in chromatin mobility that we report here, where no damage was induced, were systematically accompanied by an increase in distances between the arrays along the chromosome, a parameter which can be explained by a change in chromatin structure (Hauer et al., 2017; Amitai et al., 2017; Herbert et al., 2017). The higher order chromatin structure modification due to H2A-S19E was previously proposed to correspond to a decreased compaction (Downs et al., 2000), in agreement with a more relaxed profile of 2μ plasmid from H2A-S12E containing cells and a faster digestion of the H2A-S129E chromatin by MNase (Downs et al., 2000). The fact that in the H2A-S129E mutant, the protection pattern or the length of nucleosomal repeats was conserved (Downs et al., 2000), is also compatible with a stiffer chromatin, but the number of nucleosomes per length unit (i.e. compaction) remains to be determined. Similarly, treatment with Zeocin has been shown to cause nucleosome loss, as documented by histones H3 and H4 degradation (Hauer et al., 2017). Histone degradation could explain the additional modest effect on mobility and intra-chromosomal distances observed in H2A-S129E mutant when treated with Zeocin since the global level of H4 was not affected in undamaged H2A-S19E cells, but substantial degradation was observed in both, WT and H2A-S19E cells in the presence of Zeocin. Thus chromatin stiffening due to H2A phosphorylation and chromatin de-compaction due to histone loss could exist together in Zeocin treated cells. Only close examination of the chromatin structure by super resolution will help to determine the nature of these chromatin changes.

Our live-cell chromosome dynamics observations combined with the genetic evidence presented here suggest a novel mode of DDR regulation via an extremely efficient modification of the chromosome structure. The activation of DDR is evident after DNA damages, which restores cell cycle arrest in the double mutant Δ*rad9*, H2A-S129E. Which effectors could be responsible for the reactivation of the checkpoint? It was tempting to test whether the phosphorylation of Rad53 was restored under these conditions. The lack of detectable phosphorylated form of Rad53 indicates that downstream checkpoint activation occurs through other effectors that have yet to be identified. Another hypothesis is linked to the Mad2-mediated Spindle Assembly Checkpoint (SAC) that we previously show to sustain arrest (Dotiwala et al., 2010). Activation of this extended G2/M arrest is dependent on formation of γ-H2AX (Dotiwala et al., 2010). A possible explanation to the bypass of Δ*rad9* by H2A-S129E could be an extended arrest through the Mad2 SAC. Regardless the nature of the effectors activated by H2A-S129E, our results indicate that H2A post-translational modification of the chromatin that can be achieved in the absence of any protein synthesis, results in a very proficient DDR, probably faster than any transcriptional regulation.

### What is the function of H2A-S129E?

In the presence of H2A-S129E, Rad9 is no longer essential for arresting cells in G2/M after DNA damage or for facilitating damage-associated chromosome mobility. Based on the evidence presented here, we propose that chromosome mobility induced by chromatin structural changes, is a manner to efficiently signal damage and help repair. The chromatin changes and the enhanced dynamics may favor accessibility of the NHEJ repair machinery to the extremities of the break. In agreement with this, Tel1^ATM^ that phosphorylates H2A-S129, facilitates NHEJ repair protein recruitment and prevents dissociation of broken DNA ends (Lee et al., 2008). Consequently, inappropriate rejoining of chromatin fragments that can result in genetic translocations is decreased, as we and others have observed both in yeast and mammals (Reina-San-Martin et al., 2003; Celeste et al., 2003; Lee et al., 2008). Besides, DSB ends resection is thought to regulate the ƴ-H2A(X)-mediated recruitment of remodeling complexes, SWR1, Fun30 and INO80, to promote repair by respectively NHEJ and HR (van Attikum et al., 2007; Horigome et al., 2014; Eapen et al., 2012). Notably, the interaction between Fun30 and H2A-S129E chromatin is impaired (Eapen et al., 2012), raising the possibility that resection factors in H2A-S129E mutant would also be impaired, thus favoring NHEJ. The role of chromosome mobility in repair by NHEJ is consistent with the observation that increased roaming of deficient telomeres facilitates NHEJ in mammals (Lottersberger et al., 2015; Dimitrova et al., 2008). Concerning translocations, i.e. the junction between two DSBs on distinct chromosomes, two theories are proposed. On the one hand, the limited motion of DSB ends would have a greater probability of forming a translocation provided they are spatially close, on the other hand, mobile ends would favor the connection between ends which have lost their proper interaction (Roukos et al., 2013; Lottersberger et al., 2015). Thus a NHEJ made efficient thanks to the mobility of DSB extremities would counterbalance an ectopic repair between two distant breaks. Our results favor this last hypothesis. Mobilizing DSBs to promote faithful repair may be preserved by evolution. The reason why such H2A-S129E mutation has not been positively selected may be linked to the fact that it would be useful to keep such additional level of regulation, through increased mobility, otherwise impossible to reach in the case of “permanent” increased mobility.

Finally, the evolutionary conservation of chromosome mobility nurtures the possibility that similar mechanisms through chromatin modification exist in other organisms. In both yeast and mammals, the initial recognition of DSBs and activation of the checkpoint kinases is not dependent on the formation of γ-H2A(X) (Celeste et al., 2003; Dotiwala et al.; Kruhlak et al., 2006), but this modification is important in maintaining damage arrest and in the recruitment of DSB repair factors. In mammalian cells, the response to DNA damage is complex and includes modification due to damage include protein modifications that do not exist in yeast, such as the poly ADP-ribosylation by poly (ADP-ribosyl) polymerases (PARPs). PARP also induces chromatin mobility (Sellou et al., 2016). It would be interesting to test if chromatin mobility induced by parylation in mammalian cell lines helps to preserve chromatin integrity.

---

**Supplementary Figure 1.**
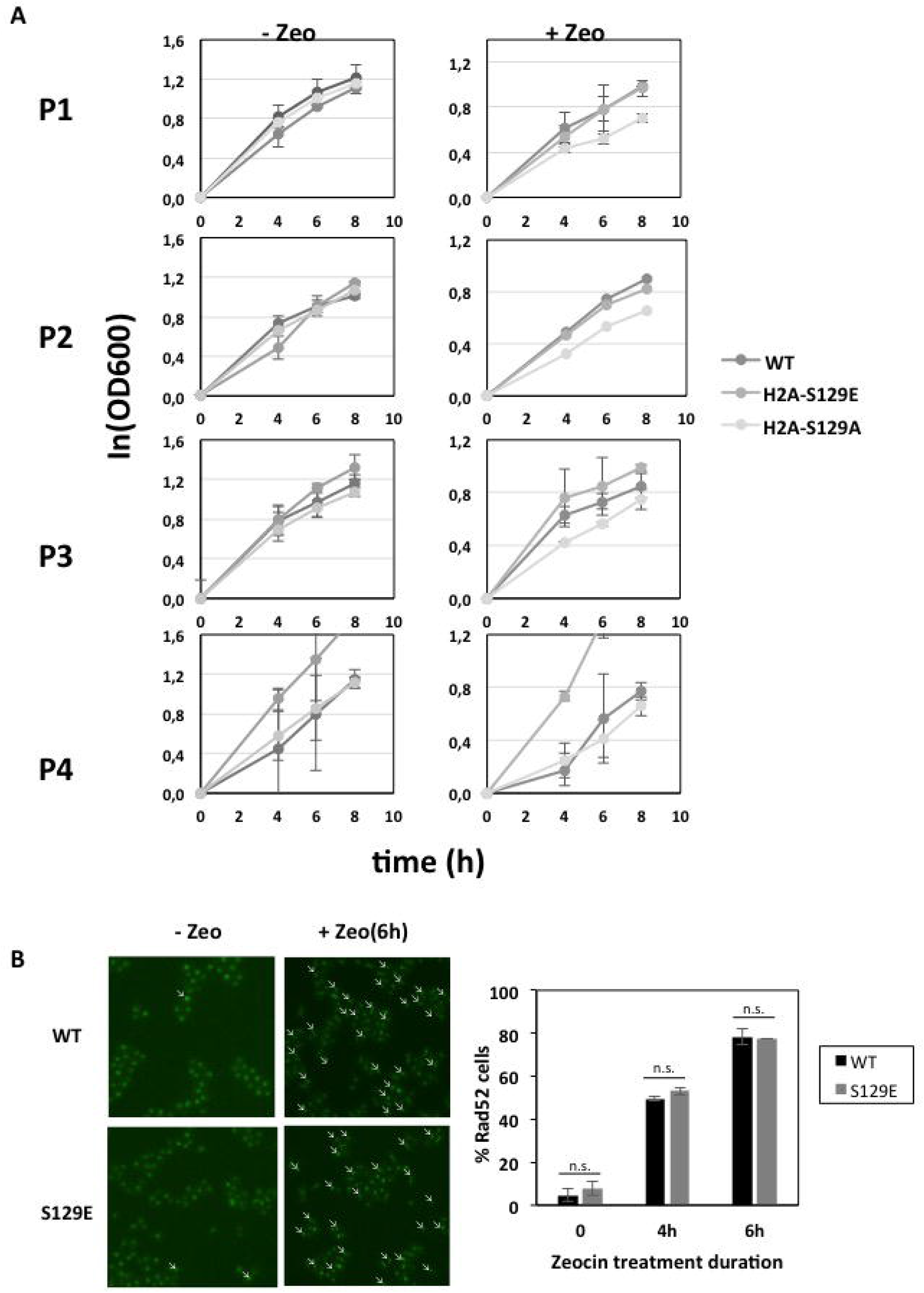
Absence of growth defects and intrinsic DNA damages in H2A-S129E mutant. A. Representative growth curves for the WT, H2A-S129A, H2AS129E strains in the absence (left) or presence (right) of 250μg/ml Zeocin treatment. Mean values for two independent experiments are plotted for each time point, with error bars showing standard error of the mean (SE). B. No intrinsic DNA damage in H2A-S129E in absence of Zeocin treatment but prolonged exposure to Zeocin increases DNA damage. Rad52-GFP foci are shown (arrowheads) in representative images of yeast cells that were either untreated or exposed to 250μg/ml of the genotoxic drug Zeocin for 4 and 6 h in WT (black) and H2A-S129E strain (grey). Bar graphs show mean ± s.e.m. P values are calculated after a non-parametric t test (n.s, not significant P>0.05).

**Supplementary Figure 2.**
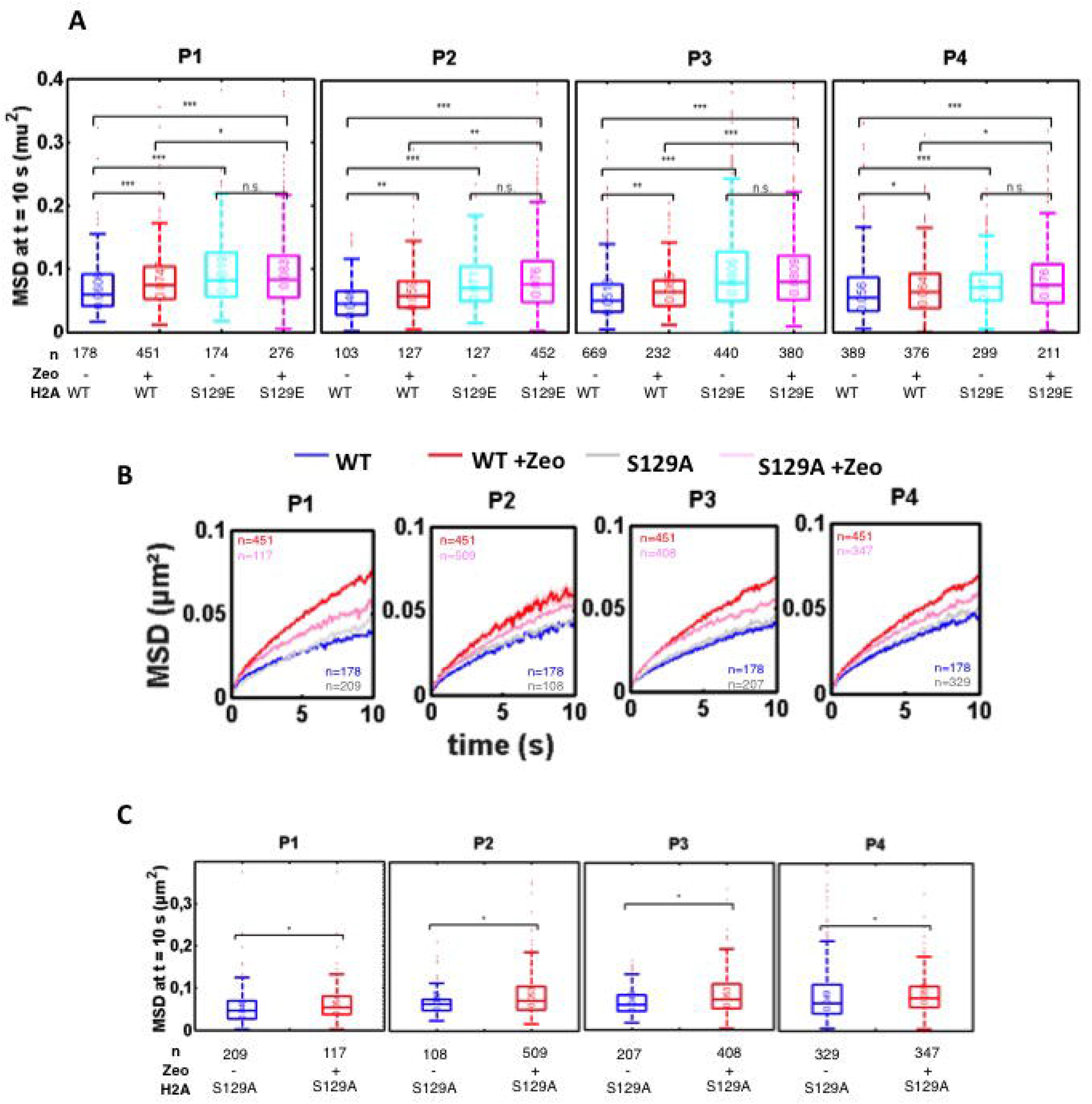
Values of MSDs at 10 sec. in WT, H2A-S129E and H2A-S19E mutated strains. **A**. Values of MSD at 10 sec for WT, H2A-S129E and H2A-S129A. Boxplots show the distribution of MSD at 10 s in absence of Zeocin (WT, blue; H2A-S129E, light blue) or after 3 h Zeocin exposure (WT, red; H2A-S129E, pink), for the four loci Gr1–Gr4. The horizontal line at the center of each box indicates the median value, the bottom and top limits indicate the lower and upper quartiles, respectively. The whiskers indicate the full range of measured values, except for outliers, which are shown as small red dots. Brackets indicate the result of a Wilcoxon rank-sum test between distributions, with “n.s.” for “not significant” (P > 0.05), * for P < 0.05, ** for P < 10^−2^ and *** for P < 10^−3^. **B**. MSDs for WT and H2A-S129A. Mean square displacement measured as in figure 1D as function of time interval of the four Gr1 to Gr4 green loci in H2A-S129A background. Blue and red curves are for untreated and treated Wild-Type cells (WT); grey and pink curves are for untreated and treated H2A-S129A mutant, respectively. The numbers of cells used to compute each curve (n) are indicated. **C**. Values of MSD at 10 sec for H2A-S129A in the presence or absence of treatment. Boxplots show the distribution of MSD at 10 s as in B, in absence of Zeocin (blue) or after 3 h Zeocin exposure (red), for the four loci Gr1–Gr4 in H2A-S129A mutated background.

**Supplementary Figure 3.**
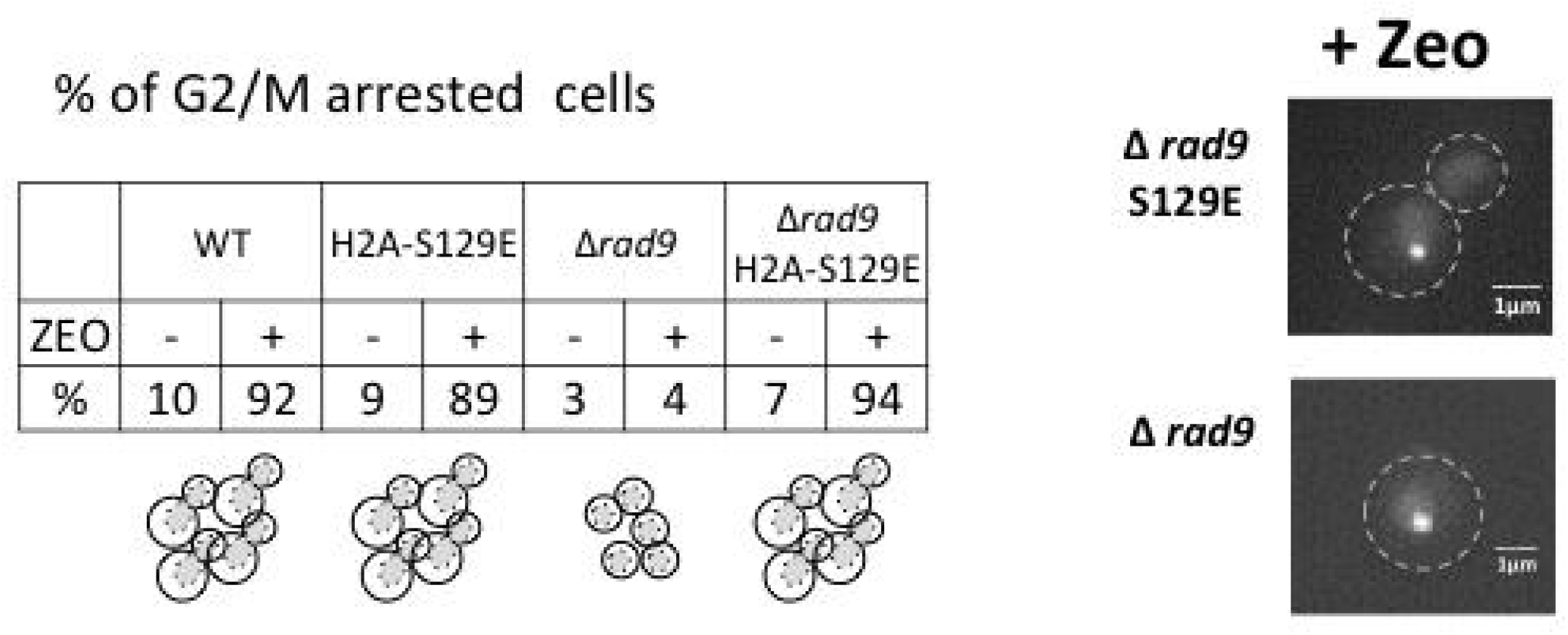
Percent of G2/M arrested cells in the absence or the presence of Zeocin for WT, H2A-S129E, Δ*rad9* and Δ*rad9* H2A-S129E mutated cells. Cells expressing fluorescent LacI-GFP protein to lacO-array were inspected after image acquisition. Unbound LacI-GFP was used as a nuclear staining. G2/M arrested cells show nuclear masses separated between mother and daughter cells, but no septum formed. Examples are shown on the right.

**Supplementary Figure 4.**
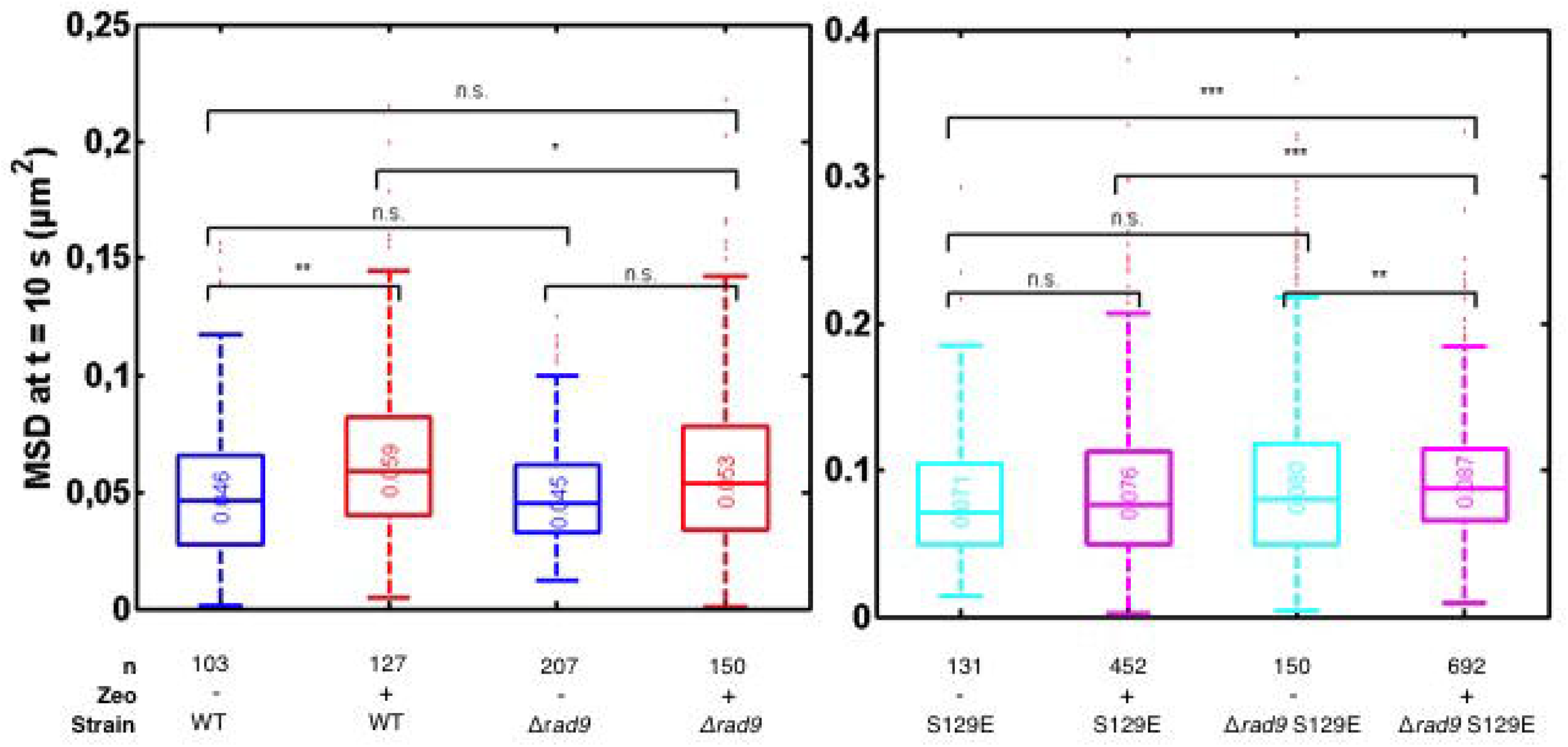
Values of MSDs at 10 sec for WT, H2A-S129E, Δ*rad9* and Δ*rad9* H2A-S129E mutated cells. Boxplots show the distribution of MSD at 10 s of WT compared to Δ*rad9* strains (left) and H2A-S129E compared to Δ*rad9* H2A-S129E double mutant (right) in absence of Zeocin (bleu and cyan, respectively) or after 6 h Zeocin exposure (red and magenta, respectively). The horizontal line at the center of each box indicates the median value, the bottom and top limits indicate the lower and upper quartiles, respectively. The whiskers indicate the full range of measured values, except for outliers, which are shown as small red dots. Brackets indicate the result of a Wilcoxon rank-sum test between distributions, with “n.s.” for “not significant” (P > 0.05), * for P < 0.05, ** for P < 10^−2^ and *** for P < 10^−3^. Analyzed cells range from (n) ~100 - ~1000.

**Supplementary Figure 5.**
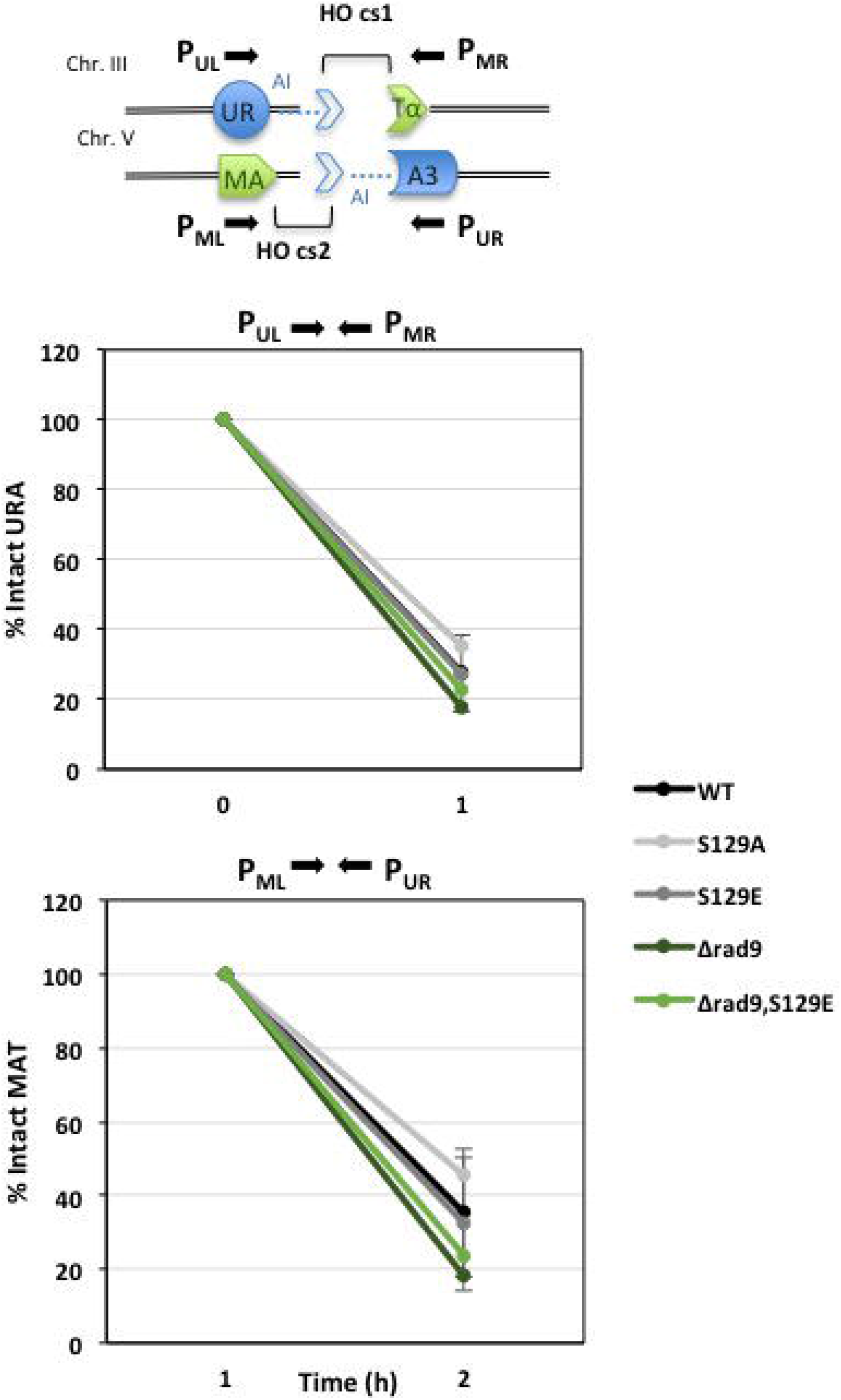
Efficiency of the two HO-cleavages in translocation assay strains by Q-PCR. Efficiency of HO cleavage in WT, H2A-S129A, H2A-S129E, *Δrad9* and *Δrad9*, H2A-S129E strains carrying the two HO cleavage sites at the MATα and *URA3* locus was determined by quantitative PCR using primers flanking HO recognition sites before (t0h) and after (t1h) galactose induction, normalized by the amount of *ACT1* sequence.

**Supplementary Table 1. Strains used in this study.**

## Materials and Methods

### Strains construction

All strains, plasmids and primers used in this study are listed in Table 1. Fluorescent labeled strains used in this study were constructed by insertion of Tet-Operator and Lac-Operator arrays inserted at specific locations on the chromosome IV (Robinett et al., 1996). These arrays are bound by Tet-Repressor and Lac-Inducer, which are fused to the fluorescent proteins mRFP and eGFP, respectively.

yH2A129A/ yH2AS129E mutants were constructed using CRISPR/Cas9 technology as described by (Anand et al., 2017). Briefly, a 20 nucleotide sequence, corresponding to the sgRNA, flanked by a NGG sequence (Cas9 PAM) was selected from the sequence of both H2A genes (*HTA1*/*HTA2*) containing the serine 129. A complementary sequence is designed to form a duplex with *Bpl*I overhangs and cloned into *Bpl*I-digested (pEF562) Yeast Cas9 plasmid (1:1). To introduce the modifications into the yeast genome, a single-stranded 80 nt donor sequence was designed, consisting of 40 nucleotides upstream and 40 nucleotides downstream of the targeted sequence and containing an alanine 129 (H2AS129A) or glutamic acid 129 (H2AS129E) codon instead of the serine 129. Yeast cells were transformed with 1 μg yeast Cas9 plasmid containing the sgRNA against either *HTA1* or *HTA2* (pEF567 and pEF568 respectively) and 2 μg donor sequence. After transformation, yCas9 plasmids were lost, and sequence modification of *HTA1* and *HTA2* genes were verified by sequencing.

### Yeast cell cultures for in vivo microscopy fluorescence observations

Yeast cells were grown in a selective medium overnight at 30°C, diluted 1:50 in the morning and grown for two generations. When relevant, Zeocin at a concentration of 250 μg/ml was added to the culture 6 h before imaging. Cells reaching exponential phase (1 O.D. at 600 nm) were then centrifuged, concentrated to 1.5 10^7^ cells/ml and 3μl were spread on agarose patches (made of synthetic complete medium containing 2% agarose). Patches were sealed using VaLaP (1/3 Vaseline, 1/3 Lanoline, and 1/3 Paraffin).

### Wide-field microscopy

Live cell imaging was done using a wide field microscopy system featuring a Nikon Ti-E body equipped with the Perfect Focus System and a 60x oil immersion objective with a numerical aperture of 1.4 (Nikon, Plan APO). We used an Andor Neo sCMOS camera, which features a large field of view of 276 × 233 μm at a pixel size of 108 nm. We acquired 3D z-stacks consisting of 35 frames with z-steps of 300 nm using a dual band filter set (eGFP, mRFP). For each z position, two color channels are consecutively acquired with an exposure time of 100 ms. The complete imaging system including camera, piezo, LEDs (SpectraX) is controlled by the NIS-elements software.

### Wide-field image analysis and statistics

Image analyses were performed using Fiji plugins (Herbert et al., 2017). Briefly, images were corrected for chromatic aberrations using the plugin “Descriptor-based registration (2d/3d)”. Once correction applied to all acquired images, the “Detect ROI” script allowed automatically selecting non-dividing cells and computing the (x, y) coordinates of the red and green loci using a custom-made Fiji plugin implementing Gaussian fitting. For tracking in time-lapse microscopy, we used the same custom-written Fiji plugin, which allowed extracting locus positions over the entire time course for each nucleus. A custom-made MATLAB script that corrected global displacements and computed MSD curves for each trajectory using non-overlapping time further analyzed these trajectories. Finally, other MATLAB scripts were used to fit power laws to individual MSD curves or population-averaged MSDs over time intervals 0.1 - 10 s.

### Colony Forming Unit assay

Each strain was grown overnight in 3 ml of YPD. The day after each culture was appropriately diluted and ~ 200 colonies were plated onto YPD plates and YPD plates supplemented with Zeocin at a concentration of 12.5 μg/ml in order to induce random DSBs (as described in Seeber et al., 2013). Colonies were counted after 2–3 days of incubation at 30°C. For each strain, three independent experiments were performed with the corresponding controls.

### Plasmid repair assay

The plasmid pRS413 (*HIS3*) was linearized with either *EcoRI*, which generates cohesive ends. Equal numbers of competent cells are transformed with 100ng of either linear or circular plasmids. Following transformation with linear plasmid, the cell must repair the *HIS3*-containing plasmid to survive subsequent plating on Dropouts plates lacking histidine. Each strain’s repair efficiency was quantified, relative to the isogenic wild type, by assessing the number of colonies obtained with the cut plasmid relative to the circular plasmid. Each mutant was assayed a minimum of three times in triplicate, the averaged results, standard errors and unpaired t tests are reported.

### Translocation assay

Strains were grown overnight in 20mL of synthetic minimum medium SC (2% raffinose) at 30°C, diluted to a density corresponding to an OD of 0.2 at 600nm, and grown for 3 h in the same synthetic minimum medium in order to reach exponential growth phase (OD = 1). Cultures were diluted appropriately to have ~10^4^ and ~10^3^ colonies/plate after counting with Malassez chamber and plated onto Galactose containing SC dishes lacking or not Uracile. Colonies were counted after 2-3 days of incubation at 30°C. Translocation efficiency (%) is done by calculating the ratio between the number of colonies in −URA SC plates containing Galactose and the number of colonies in non selective SC plates containing Galactose. For each strain, three independent experiments were performed.

### FACS analyses

Cells were grown to mid-log-phase in liquid cultures, and treated or not with Zeocin at 250 mg/μl during 6h at 30°C. After incubation, samples were fixed in 70% ethanol and kept at 4°C for 48 h. Cells were then resuspended in 50 mM Sodium Citrate (pH 7) containing RnaseA at 0.2 mg/mL final concentration. After incubation at 37°C for 1h, Sytox Green was added to a final concentration of 1mM. A total of 10^6^ cells were analyzed with a CANTO II flow cytometer (BD Biosciences). Aggregates and dead cells were gated out, and percentages of cells with 1C and 2C DNA content were calculated using FLOWJO software.

### Western Blotting

Cells were grown to mid-log-phase in liquid cultures, and then treated with Zeocin at 250 mg/μl during 3h at 30°C. After incubation, samples were washed once with water and resuspended in 20% TCA. Cells were lysed by sonication three times for 30 sec. and the protein lysates were pelleted by centrifugation at 14000 rpm for 5 min at 4°C. The pellets were dissolved in 1x SDS sample loading buffer by boiling for 5 min. Samples were centrifuged for 30 sec. at 18000 g in a microcentrifuge and the supernatant was retained as the protein extract. Protein samples were resolved on a Bolt™ 10% Bis-Tris Plus gel and transferred onto a polyvinylidene difluoride membrane (Immobilon-P; Millipore). Membranes were probed with anti-H4 (ab10158) and anti-HA-HRP (26183-HRP) antibodies. As a loading reference, we use anti-Dpm1 (5C5A7) anti-Actin antobody (MA1-744). Anti-mouse and anti-rabbit IgG HRP-conjugated secondary antibodies were obtained from Thermofisher (A11005 and A31556). Blots were developed using the ECL plusWestern Blotting System (GE Healthcare).

## Acknowledgments

We thank S. Eun Lee and K. Dubrana for strains and advice, S. Duchez for her help with the FACS experiments and A. Carré Simons for drop tests. We acknowledge P. Lesage, A. Bonnet, P. Therizols, A. Canat and D. Waterman for their helpful comments on the manuscript. The authors would also like to thank their respective team members for very fruitful discussions.

